# The common oral parasitic protozoan *Entamoeba gingivalis* forms cysts in response to antibiotic treatment

**DOI:** 10.1101/2022.12.19.521153

**Authors:** Christin Becker, Aysegül Adam, Henrik Dommisch, Thomas Stach, Arne S. Schaefer

**Author notes:** Corresponding author, Arne S. Schaefer, **Email:**. **Competing Interest Statement:** The authors have no competing interests to disclose.

## Abstract

The eukaryotic protozoan *Entamoeba gingivalis* (*E.g*.) is strongly associated with inflamed periodontal pockets. Unlike other obligate anaerobic *Entamoeba* species, it is considered to not have a life cycle of actively dividing trophozoites and dormant cysts. Accordingly, it has been regarded as non-infectious. To investigate if *E.g* is capable of encystation in response to adverse environmental conditions, we cultivated clinical isolates of *E.g*. collected from inflamed periodontal pockets in antibiotics for 8 days. The cytomorphological and ultrastructure forms of the amoeba were investigated by transmission and scanning electron microscopy to reveal cyst formation. We observed exocysts and the encapsulated trophozoids separated by an intra-cystic space, a dense poorly vesiculated cytoplasma and polygonal surface areas of cysts. The cysts walls were composed of chitin. Cysts were conspicuously smaller compared to trophozoids and lacked pseudo- and filipodia. We did not observe multi-nucleated trophozoids after antibiotic induces encystation. Cyst formation in *E.g* may explain why established treatment approaches often do not stop periodontal tissue destruction during periodontitis and periimplantitis.

## Introduction

Periodontitis is a very common complex inflammatory disease of the oral cavity with a frequency of 47% for adults >30 years of age in Western countries [1–3]. It is characterized by gingival bleeding caused by active inflammation in conjunction with progressive destruction of the tooth-supporting apparatus including recession of the gingiva and resorption of the alveolar bone, leading to tooth loss. Necrotizing ulceration of the interdental papilla and the periodontal and alveolar ligament also characterize some forms of periodontitis [4].

Periodontitis is a major public health problem. This is, because the long-lasting inflammation of the oral mucosa has a negative impact on general health and periodontal infections are associated with a range of systemic diseases leading to premature death, including cardiovascular diseases and diabetes [5]. Additionally, since it leads to tooth loss and disability, if untreated, periodontitis significantly impairs quality of life, results in significant dental care costs and is a source of social inequality. It results from a polymicrobial insult, including as yet unidentified and never cultivated bacterial species and possibly viruses, fungi, and eukaryotic parasites [6].

The anaerobic protozoan parasite *Entamoeba gingivalis* (*E.g*.) is observed in inflamed periodontal pockets with a prevalence of > 80 % [7, 8]. It has a strong pathogenic potential, which is implied by the observations that *E.g*. is able to invade lacerated oral mucosa where it kills oral epithelial cells by trogocytosis, a process in which it physically extracts and ingests cellular material from human epithelial cells. In appearance, this process is similar to trogocytosis of colon epithelial cells by *Entamoeba histolytica* (*E.h*.), a human anaerobic pathogenic *Entamoeba* species that colonizes the human gut and is the causal parasite of amebic dysentery. Similarly to infection of colon epithelium with *E.h*., infection of oral epithelial cells with *E.g*. causes significant induction of proinflammatory cytokine signaling cascades, leading to >1500 fold increased expression of interleukin-8 [9], [7, 10], which is the major cytokine for attraction and activation of neutrophils in inflammatory regions.

Despite the shared pathogenic potential, its considerable abundance in bleeding periodontal pockets and the significant association with severe forms of periodontitis, to date, *E.g*. is being considered commensal and not infectious [11]. A significant argument for this consideration is that essential for infection and the epidemiological impact of *E.h*. is the capability to form cyst walls. These structures are composed of chitin, which are hard and impermeable to small molecules and protect the parasite from natural environmental insults outside of the colon, such as environmental oxygen, drying, osmotic shock in water, or lysis by stomach acids and duodenal proteases. Due to their wall organization with a high content of chitin, amoeba cysts also confer resistance to various antibiotic medication and therefore represent a serious problem in the treatment of amoebic infections. In contrast to *E.h*. and other anaerobic *Entamoeba* species, cyst formation of *E.g*. has not been observed. Accordingly, it is currently thought that E.g., which is a facultative anaerobic protozoon, is not able to form cysts and therefore not infectious. Accordingly, demonstrating that *E.g*. is able actively responds to adverse environmental conditions by the formation of cysts would gives significant evidence for a serious potential of this parasite for infection and resistance, and therefore affecting therapeutical approaches and our understanding of the etiology of periodontal diseases.

We hypothesized that fine structural organization of *E.g*. after exposure to an unfavorable environment can determine *E.g*.’s potential to form cysts. To test our hypothesis, we examined *E.g*.’s cytomorphological features, ultrastructure forms and biochemical characteristics after antibiotic treatment of *E.g*., cultured *in vitro* from inflamed periodontal pockets of periodontitis patients. For this purpose, we used transmission (TEM) and raster electron microscopy (SEM) as well as fluorescence and light microscopy. We found numerous cellular structures that generally characterize amoebic cysts.

## Results

We cultivated *E.g*. from dental plaque of inflamed periodontal pockets of periodontitis patients under anoxic conditions. After 2 days of cultivation, *E.g*. trophozoites showed their characteristic mobile forms (**Figure 1**). The cytoplasma contained multiple food vacuoles and a contractile vacuole. Peripheral chromatin characterized the single nucleus. TEM additionally showed the characteristic circular karyosome with a central body and clusters of small round shaped electron dense nuclear bodies. The trophozoites generally presented a longitudinal shape continuously forming one or more pseudopods at different positions during locomotion. REM showed the presence of various pseudopods and filipodes, thin filaments of short and elongated forms. The lengths of the trophozoids varied in a range of 10-20 μm in diameter.

**Figure 1.**
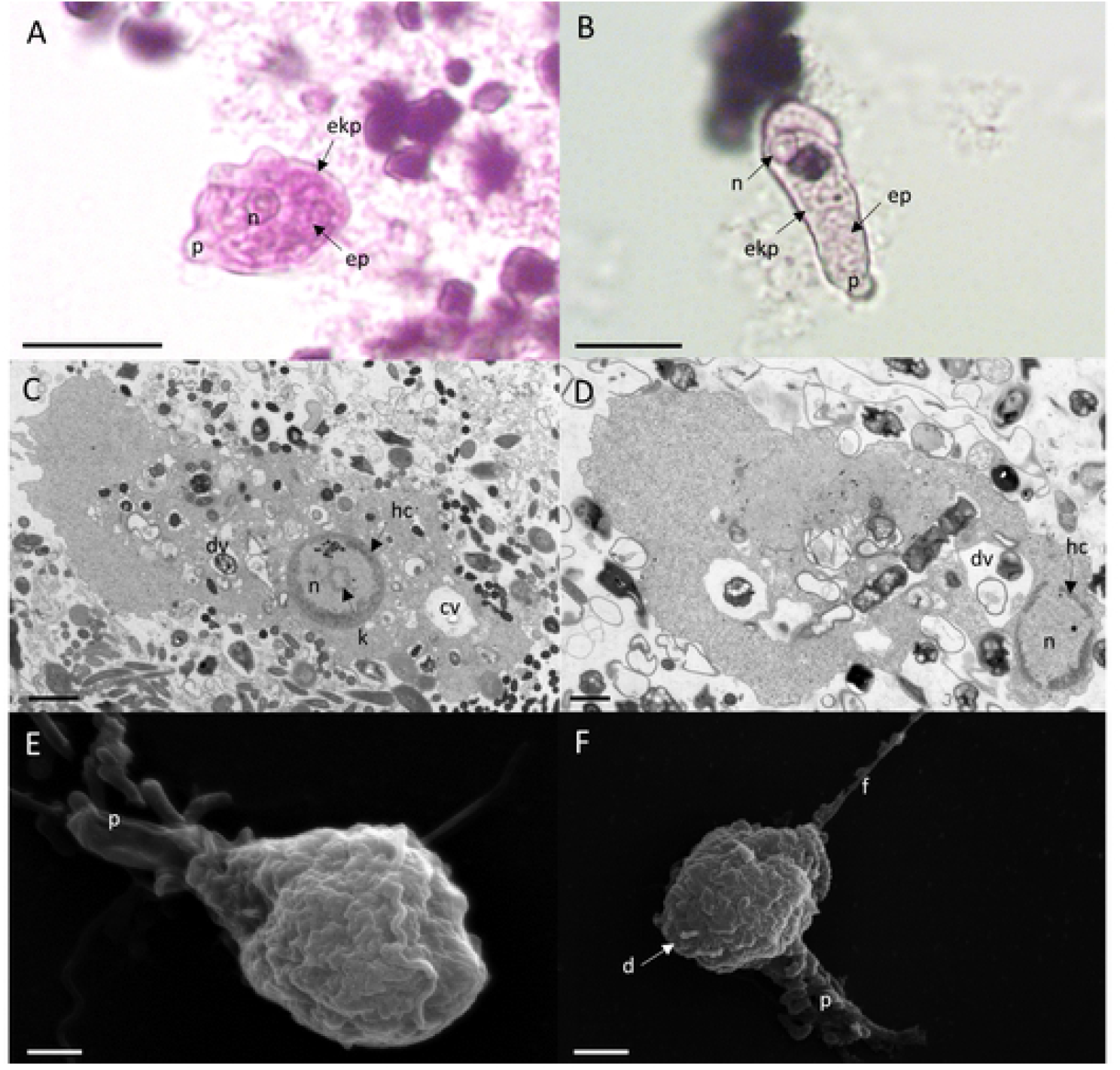
Cytomorphological features and ultrastructure forms of *E.g*. trophozoites. Upper panel: Trophozoites under the light microscope show a distinct nucleus (**n**), pseudopodia (**p**), a granoulous endoplasma (**ep**) and a transparent ektoplasma (**ekp**), which locates adjacent to the distinct plasma membrane as characteristic features (A, B). Scale bar = 10 μm Middle panel: TEM shows the characteristic features of the nucleus (**n**) and cytoplasma of the trophozoite. The nucleus has a circular karyosome (**k**), condensed peripheral heterochromatin (**hc**) and additionally shows a cluster of small round shaped electron dense nuclear bodies. In the cytoplasm, numerous digestive vacuoles (**dv**) and a contractile vacuole (**cv**) are visible (C, D). Scale bar = 1 μm Lower panel: REM shows surface ultrastructures of trophozoites that are characterized by a granulose wrinkled membrane and smooth pseudopods (**p**), filipodes (**f**), and digipodes (**d**) (E, F). Scale bar = 2 μm

After 2 days, the culture medium was replaced with fresh medium added with antibiotics amoxicillin and metronidazole and the cultures kept on growing for 8 days. In cultures that were treated for 8 days in medium containing the antibiotics amoxicillin and metronidazole, we could observe trophozoite cells of *E.g*. with typical ultrastructural features of amoebic cysts (**Figure 2**). With a diameter of 3-6 μm cysts were conspicuously smaller than trophozoids. The cytoplasm was poorly vesiculated and appeared less condensed in comparison with the cytoplasm of unencapsulated trophozoids, which in contrast contained numerous vesicles, revealed by TEM. Some cysts also showed a large, more electron transparent compartment that may contain glycogen as described for other *Entamoeba* species [12]. The plasma membrane of the encysted trophozoids was wrinkled and an electron-transparent intra-cystic space clearly separated the trophozoid from the cyst wall, with few flanking points where the plasma membrane and the inner side of the cyst wall were closely apposed. In some cysts, these flanking points showed numerous, perpendicular thin filaments that connected the plasma membrane with the cyst wall, possibly at the position of putative ostioles, small openings through which the trophozoid might escape from the cyst (**Figure 3**). These complexes were covered by flat structures, possibly serving as operculae to close the apertures of the cyst when the soft parts of the trophozoids are retracted. Within the intra-cystic space at the area where the cytoplasmic membrane adhered to the exocyst, in some cells, we observed vesicles of 8-10 nm diameter, which fused with the plasma membrane and the exocyst. This process indicated a growing exocyst and incomplete cyst formation. Additionally, in some cells we noted the presence of putative splintered chromatoid bodies. For several *Entamoeba* species, chromatoid bodies have been described to arise in early cystic stages by aggregation of ribosomes forming dense crystalline structures in the cytoplasm, which in maturing amoebic cysts fragment into separate particles [13]. SEM revealed that *E.g*. cysts were characterized by pronounced polygonal edges at the exterior surface. These edges clearly converged at the putative osteole and operculum.

**Figure 2.**
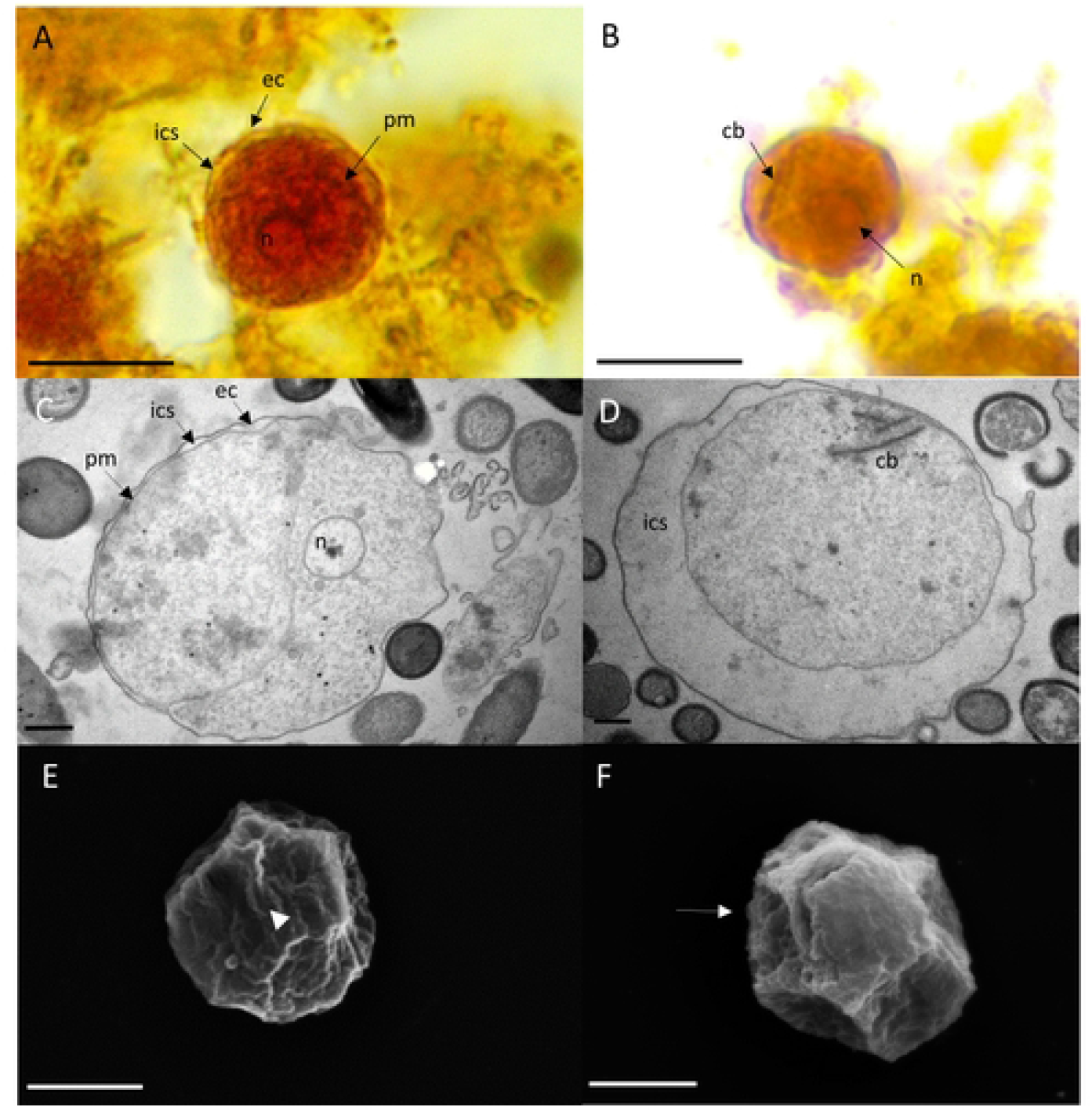
Cytomorphological features and ultrastructure forms of *E.g*. cysts. Upper panel: Encysted trophozoids are visible under the light microscope with the wrinkled plasma membrane (**pm**) being separated from the encasing exocyst (**ec**) by a clear intercystic space (**ics**). The trophozoite has a single nucleus (**n**) (A). In some cells, a chromatoid body (**cb**) is visible outside of the nucleus (B). Scale bar = 10 μm Middle panel: TEM reveals characteristic features of mature cysts. The trophozoite’s plasma membrane is separated from the exocyst by an intra cystic space. The cytoplasma is homogeneous and does not contain digestive vacuoles. A compartment is more electron transparent and may contain glycogen (C). The encysted trophozoite has a single nucleus and splintered chromatin bodies (D). Scale bar = 1 μm **Lower panel**: Ultrastructure of polygonal cysts with a distinct ostiole (**arrowhead**) (E) and operculum (**arrow**) (F). Scale bar = 2 μm

**Figure 3.**
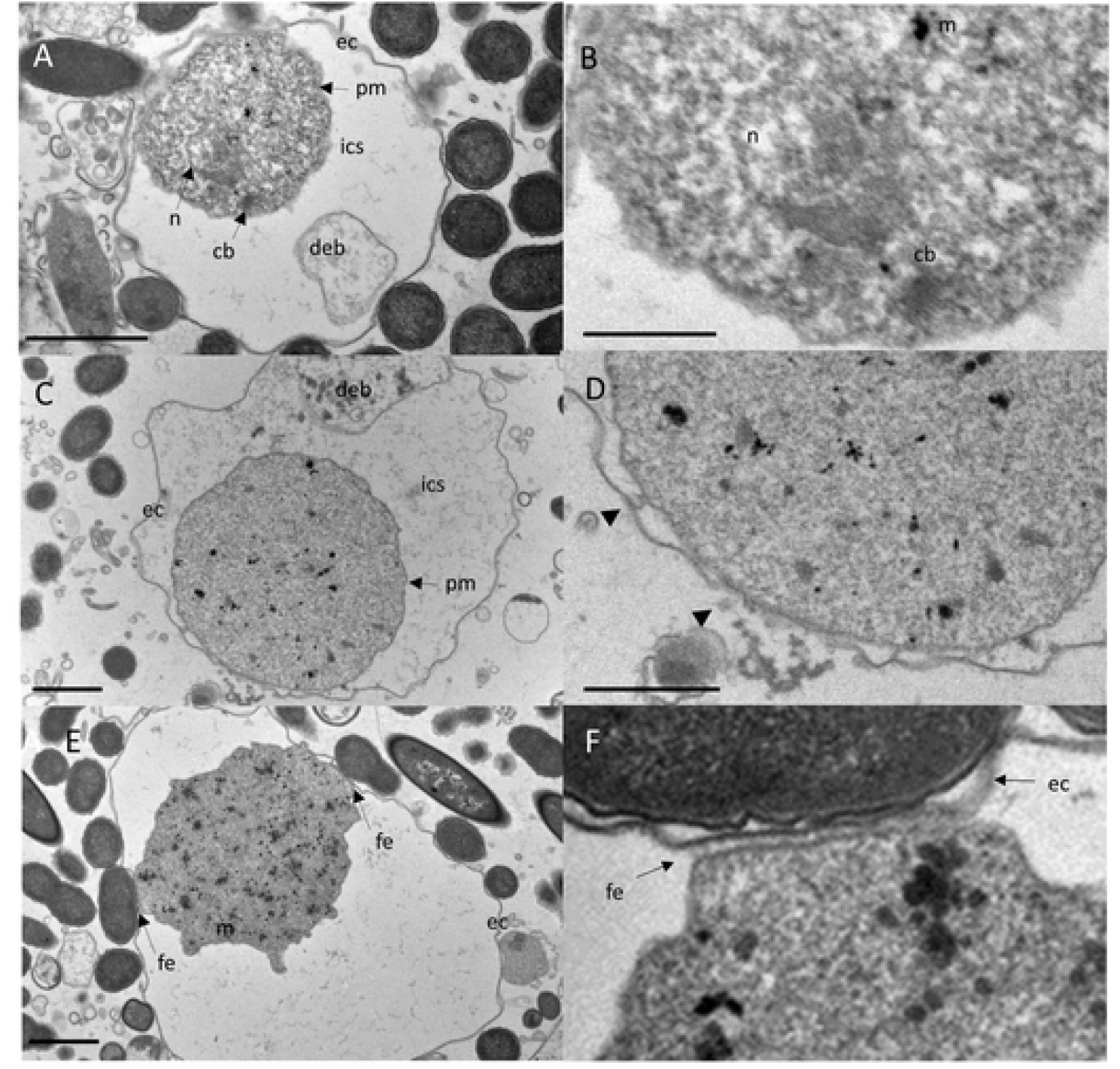
Details of cytomorphological features of the mature cysts revealed by TEM. Upper panel: The trophozoite’s plasma membrane is separated from the exocyst by an electron-transparent intra cystic space (**ics**). Cell debris (**deb**) is encased within the ics (A). Close-up view of A highlights an electron-dense capped mitosomes (**m**) and parallel electron-dense structures of a putative truncated chromatin body (**cb**) adjacent to the nucleus (B). Scale bar = 500nm Middle panel: A cyst developing the exocystic wall (C). Close-up view of C, showing ec close at the pm with several vesicles free and fused with pm and ec (**arrow heads**), indicating a growing cyst wall and incompletion of encystment. The cyst wall is developing bidirectional in close proximity to the trophozoite (D). Scale bar = 500nm Lower panel: An encysted trophozoid with two flanking points where pm is closely apposed to ec (E). Close-up view of C, showing the connection of the trophozoid to the cyst wall, which consists of an array of perpendicular fibrillar elements (**fe**) (F). Scale bar = 1μm

A major scaffolding component of the cyst wall of *Entamoeba* species like *Entamoeba histolytica, Entamoeba dispar* and the reptilian parasite *Entamoeba invadens* is chitin, a homopolymer of beta-(1,4)-linked N-acetyl-D-glucosamine [14]. To visualize chitin at the whole cyst level and to give direct evidence that the cyst wall like structures observed by TEM and SEM have biochemical characteristics of *Entamoeba* cyst walls, we performed chitin staining using the common polysaccharide marker calcofluor white. We found that calcofluor white fluorescence reflected chitin content in these walls (**Figure 4**). These data give evidence that *E.g*. is capable to develop cysts after exposure to antibiotics.

**Figure 4.**
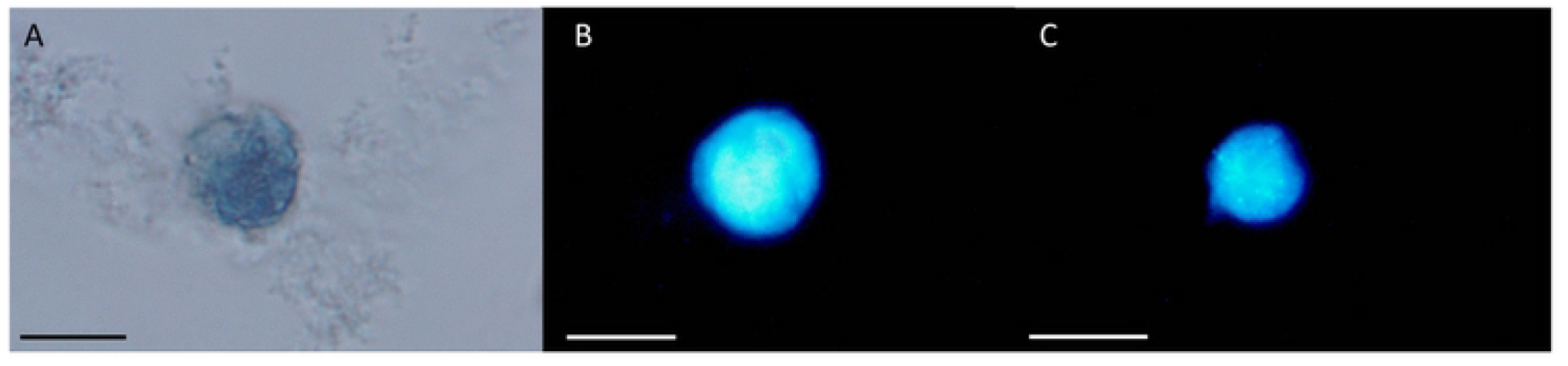
Light and fluorescence microscopy of *Entamoeba gingivalis* cysts *E.g*. cyst stained with Lugol’s iodine and calcofluor white. The polysaccharide fluorescence marker reflected chitin content in the cell wall. *E.g*. cyst under the light microscopy without UV (A). The cyst wall shows blue fluorescence under UV, indicating the presence of chititn (B, C). Scale bar = 10 μm

Finally, we observed that 10 days old *Entamoeba* cultures that were not treated with antibiotics, also differed in shape from the active trophozoites, which we generally found in 2 days old cultures. Untreated older trophozoites were characterized by a rounded shape (**Figure 5**). Light microscopy showed that the plasma membranes had irregular contact to a putative cell wall, which was occasionally separated by a transparent space. Furthermore, in these cells from 10 days old cultures, we did not observe pseudopods and filamentous structures such as digipodes and filipodes. SEM additionally revealed more structured, smoother outer surfaces compared to the outer membrane surfaces of trophozoites from 2 days old cultures. However, they lacked the polygonal edges and osteole like structures observed in cultures treated with antibiotics for 10 days. Moreover, unlike that seen in antibiotic treated 10 days old cultures, TEM revealed that the cytoplasma of untreated amoeba of the same culture age was heterogeneous and it contained, similar to trophozoites from young cultures, numerous digestive vacuoles. These characteristics implied that the amoeba in 10 days old untreated cultures were not mature cysts.

**Figure 5.**
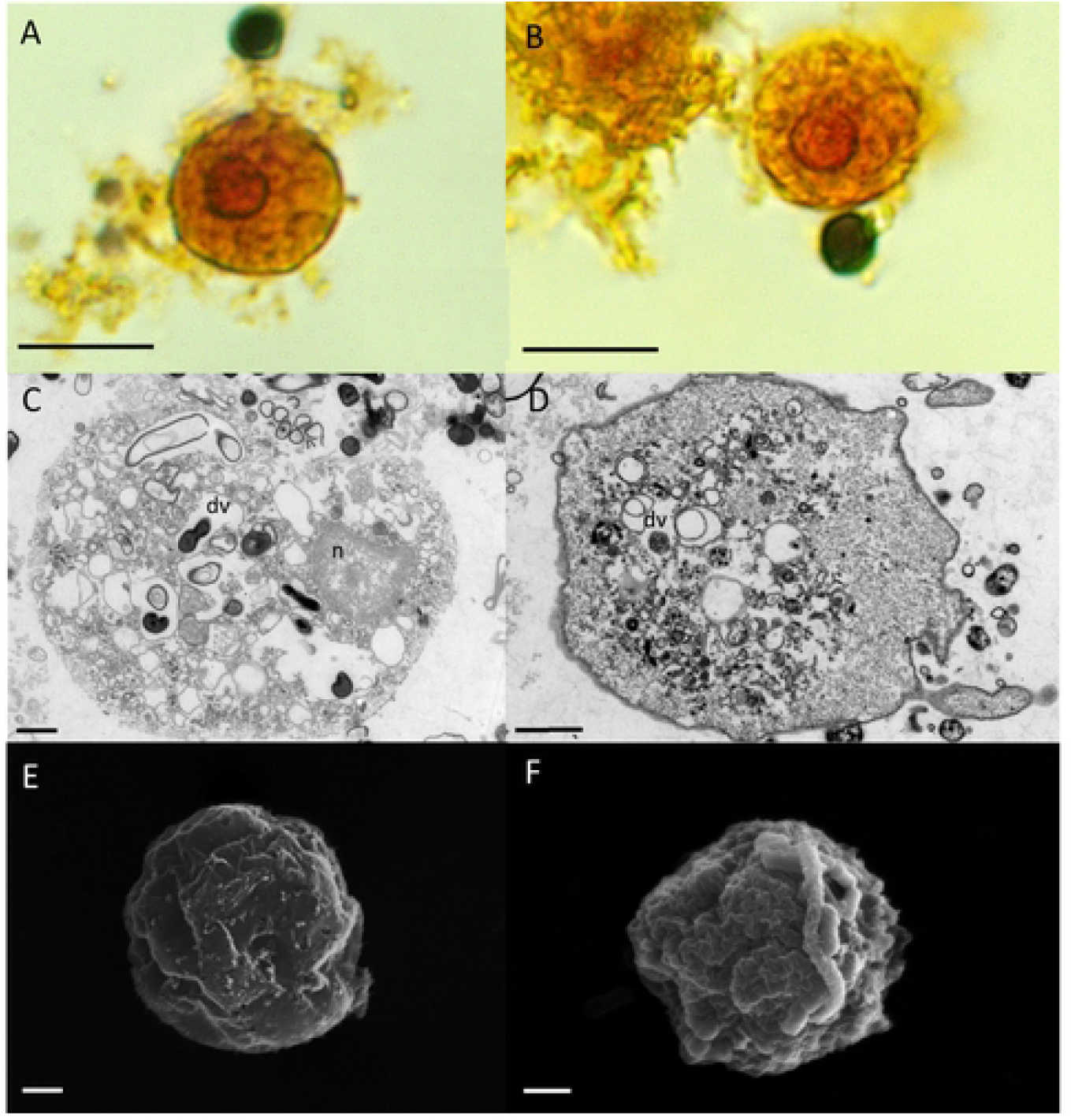
Cytomorphological features and ultrastructure forms of *E.g*. cultured for 10 days in antibiotic free axenic medium. Upper panel: Trophozoites are round shaped and show no pseudopods. Under the light microscope, the granulous cytoplasma and the nucleus with peripheral chromatin and a central karyosome is visible (A, B). Scale bar = 10 μm Middle panel: TEM reveals numerous digestive vacuoles (dv) in the cytoplasma of the round shaped trophozoites (C, D). Scale bar = 1 μm Lower panel: REM reveals a wrinkled unstructured surface of the round shaped trophozoites with neither pseuodops nor digipods and filipods (E, F). Scale bar = 1 μm

## Discussion

Using antibiotics in combination and concentrations identical to prescribed antibiotic medication in periodontal diseases, we induced cyst formation in *E.g*. trophozoides. During exposure to antibiotics, the trophozoids became smaller, changed their form to a sphere containing dense vacuole-free cytoplasma, which eventually became fully encapsulated with an exocyst. The cyst walls consisted of chitin and the plasma membranes of the spherical trophozoits were conspicuously separated by inter-cystic spaces.

This is the first time that cyst stages were observed in *E.g*. The reason for this could be that E.g. does not form cysts in inflamed periodontal pockets as their preferred natural habitat. Likewise, *E.g*. does not readily encyst in xenic cultures. Moreover, unfavorable culture conditions such as sudden and extended exposure to toxic oxygen after removal from anaerobic periodontal pockets may not allow sufficient time for protective encystation resulting in loss of the *E.g*. cultures. Perhaps this also partly explains the generally observed difficulties in culturing *E.g*. trophozoids from clinical samples.

Although we observed numerous characteristics of amoebic cyst, we did not observe multinucleated cysts, which were reported as part of the natural life cycle of *E.h*. In the natural environment, nuclear division of *E.h*. during encystation takes place in the absence of cytokinesis and the single nucleated trophozoite transforms into a multi-nucleated dormant cyst. Therefore, the function of the mature cysts outside of the host is to enable dissemination of the amoeba within an aerobic environment, which is hostile to life of *E.h*.. Therefore, development of a multi-nucleated dormant cyst is regarded as part of cell proliferation during the life cycle of *E.h*., which is paused during encystation. Whereas the multi nucleated cysts are formed for dissemination outside the human host in a hostile environment, the pre-formed multiple nuclei allow rapid completion of cytokinesis, when the amoeba recovered a favorable environment in a new host.

We did not observe multinucleated cysts. This could mean that this characteristic is not part of the life cycle of *E.g*. However, in our experiments, exposure of *E.g*. to noxious antibiotics induced cyst formation. Therefore, it is possible that the trophozoits built the observed cysts for protection instead of proliferation. Likewise, cysts of *E.h*. also show different nuclei numbers after *in vitro* induction [15] and also *in vivo* [16]. Future studies are required to determine if *E.g*. is able to form cysts as part of a reproductive cycle.

In the clinical context, cysts entail serious difficulties for the elimination of *E.g*., especially if it invaded deep periodontal pockets or gingival tissue. It may also relate to the current epidemic of periimplantitis, a destructive inflammatory process that is triggered by the presence of dental implants [17], and during which gingival tissues surrounding the implant become inflamed and subsequent alveolar bone loss over time. Cyst formation may also explain why in periodontitis or periimplantitis patients established treatment approaches fail to stop inflammatory periodontal tissue destruction. Conversion into dormant cysts if conditions become unfavorable may allow the amoeba to outlast therapeutically measures such as antibiotic treatment. However, this would require subsequent excystation of the trophozyte, a process that has already been described for *Entamoeba invadens* cysts [18], but which was not addressed in the current study.

In conclusion, the current study gives evidence that the oral eukaryotic parasite *E.g*. is able of encystation after exposure to periodontal antibiotic medication. This ability may enable this pathogen to outlast periodontal treatment and to re-colonize periodontal or implant pockets. Encystation may also have implications for dissemination and inter-individual infections. In the future, elucidation of processes involved in cyst formation may contribute to developing new treatment approaches of periodontitis, e.g. by silencing the encystation pathway.

## Materials and Methods

Subgingival plaque samples were collected from inflamed periodontal pockets of periodontitis at the Department of Periodontology, Oral Medicine and Oral Surgery, Charité - University Medicine, Berlin. The local ethics commitee approved to the study (EA1/169/20). The samples were cultured with TYGM-9 medium, covered by Nujol mineral oil, under anaerobic conditions at 35°C. After 48h, the presence of *E.g*. was examined with light microscopy and by PCR [7]. Subsequently, cultures were treated with amoxicillin (13.7 μg/ml) and metronidazole (2.5 μg/ml) for 8 days as described in [19]. **Light and UV microscopy.** For light microscopy, *E.g*. were fixated with 2.5% Glutaraldehyde for 30 min and stained with Lugol’s iodine (Carl Roth). The morphological structures of *E.g*. were analyzed under light microscopy (Leica DM750 with FlexaCam C3). For UV microscopy *E.g*. were fixed and washed three times with PBS. After centrifugation, *E.g*. were placed onto microscope slides and stained with calcofluor white M2R (Sigma-Aldrich), a fluorescent dye with specific binding to components of chitin. Fluorescence microscopy was performed with the laser scanning confocal microscope LSM 700 (Zeiss). **TEM and SEM.** *E.g*. were fixated and dehydrated as described in [20]. **TEM.** *E.g*. were embedded with propylene oxide and araldite A, stepwise increasing araldite A until 100% araldite A immersion. Araldite B was filled in to the embedding form at 60°C to achieve a bottom layer, *E.g*. were placed and left to harden. Ultrathin sections (60 nm) were cut, placed on single slot grids and contrasted with uranyl acetate and lead citrate in an automatic stainer. Ultrathin sections were examined at 50 to 80 kV in a LEO 900 transmission electron microscope at 5000x to 12000x (Zeiss). **SEM.** *E.g*. were postdehydrated with HMDS for 30 min and fixed on poly-L-lysine coverslips described in [20], mounted on aluminium stubs and then were sputter coated with 15 nm of gold in a SCD 005 sputter coater. *E.g*. were examined at 10 to 20kV in a LEO 1430 scanning electron microscope (Zeiss).

## Acknowledgments

This work was funded with a research grant of the German Research foundation DFG to AS (SCHA 1582/7-1)

